# Variability is actively regulated in speech

**DOI:** 10.1101/2021.10.08.462639

**Authors:** Ding-lan Tang, Benjamin Parrell, Caroline A. Niziolek

## Abstract

Although movement variability is often attributed to unwanted noise in the motor system, recent work has demonstrated that variability may be actively controlled. To date, research on regulation of motor variability has relied on relatively simple, laboratory-specific reaching tasks. It is not clear how these results translate to complex, well-practiced and real-world tasks. Here, we test how variability is regulated during speech production, a complex, highly over-practiced and natural motor behavior that relies on auditory and somatosensory feedback. Specifically, in a series of four experiments, we assessed the effects of auditory feedback manipulations that modulate perceived speech variability, shifting every production either towards (*inward-pushing*) or away from (*outward-pushing*) the center of the distribution for each vowel. Participants exposed to the *inward-pushing* perturbation (Experiment 1) increased produced variability while the perturbation was applied as well as after it was removed. Unexpectedly, the *outward-pushing* perturbation (Experiment 2) also increased produced variability during exposure, but variability returned to near baseline levels when the perturbation was removed. Outward-pushing perturbations failed to reduce participants’ produced variability both with larger perturbation magnitude (Experiment 3) or after their variability had increased above baseline levels as a result of the inward-pushing perturbation (Experiment 4). Simulations of the applied perturbations using a state space model of motor behavior suggest that the increases in produced variability in response to the two types of perturbations may arise through distinct mechanisms: an increase in controlled variability in response to the inward-pushing perturbation, and an increase in sensitivity to auditory errors in response to the outward-pushing perturbation. Together, these results suggest that motor variability is actively regulated even in complex and well-practiced behaviors, such as speech.

## Introduction

No matter how hard we practice, it is virtually impossible to generate exactly the same movement twice. Such variation in performance across repetitions of the same movements, or *motor variability*, is widely believed to be an inevitable consequence of noise in the nervous system, arising from stochastic events presented across all scales of brain activity, from the single-cell level to complex network dynamics (Churchland et al., 2006; Faisal et al., 2008; Renart & Machens, 2014). Indeed, many current theories of motor behavior rely on this assumption and posit that the motor system aims to minimize the detrimental effects of ‘motor noise’ on motor task performance (Harris & Wolpert, 1998; Scholz & Schöner, 1999; Todorov, 2004).

However, recent work in reaching has demonstrated that variability is not always treated as unwanted “noise” to be reduced by the motor system but may be more actively controlled. Repeated exposure to position- or velocity-dependent force fields during reaching has been shown to selectively increase task-relevant variability, potentially to facilitate more efficient future learning (Wu et al., 2014). Conversely, task-relevant variability can also be reduced when needed in some behaviors: participants exposed to a visual perturbation that magnified the horizontal displacement of the hand away from the midline during point-to-point reaching movements reduced their variability in this dimension (Wong et al., 2009). These results suggest that variability is not simply noise but can also be an important part of the signal itself that controls the motor movement (Stein et al., 2005).

Although variability poses a fundamental problem for motor control, studies on regulation of motor variability are comparatively sparse and, to date, have relied principally on relatively simple, laboratory-specific planar arm reaching tasks. While such tasks play an important role in probing motor control systems, they tackle a relatively restricted range of motor tasks that do not fully capture the complex demands of real-world behavior and, thus, may overlook some aspects of motor control in real-world tasks which are naturally less constrained. Speech production is one such task: as opposed to laboratory-specific arm reaching tasks, which involve an interaction with an uncommon external device (e.g. a joystick) and typically constrain movements to two joints, speaking is a highly over-practiced behavior that involves the coordination of roughly 100 muscles to precisely control the respiratory and phonatory systems as well as the movements of lips, jaw, velum and tongue (a muscular hydrostat with highly complex control).

Here, we aimed to test how variability is regulated in speech production, a complex, well-practiced task controlled via non-visual sensory feedback. Previous work has shown that speakers are sensitive to real-time alterations to their auditory feedback (Houde & Jordan, 1998; Purcell & Munhall, 2006). For example, speakers learn to alter their speech to oppose auditory perturbations that shift the frequencies of vowel formants (resonances of the vocal tract which serve to distinguish between different vowels). However, these studies have typically examined formant shifts that are applied in a consistent direction, regardless of the produced vowel formants, with the consequence that the mean formant values participants hear are altered, while their formant variability remains unchanged. Here, we implement a novel auditory perturbation that shifts vowel formants in a non-uniform manner, such that the mean formant values remain unchanged while trial-to-trial variability is either increased or decreased (Fig. 1).

**Figure 1.**
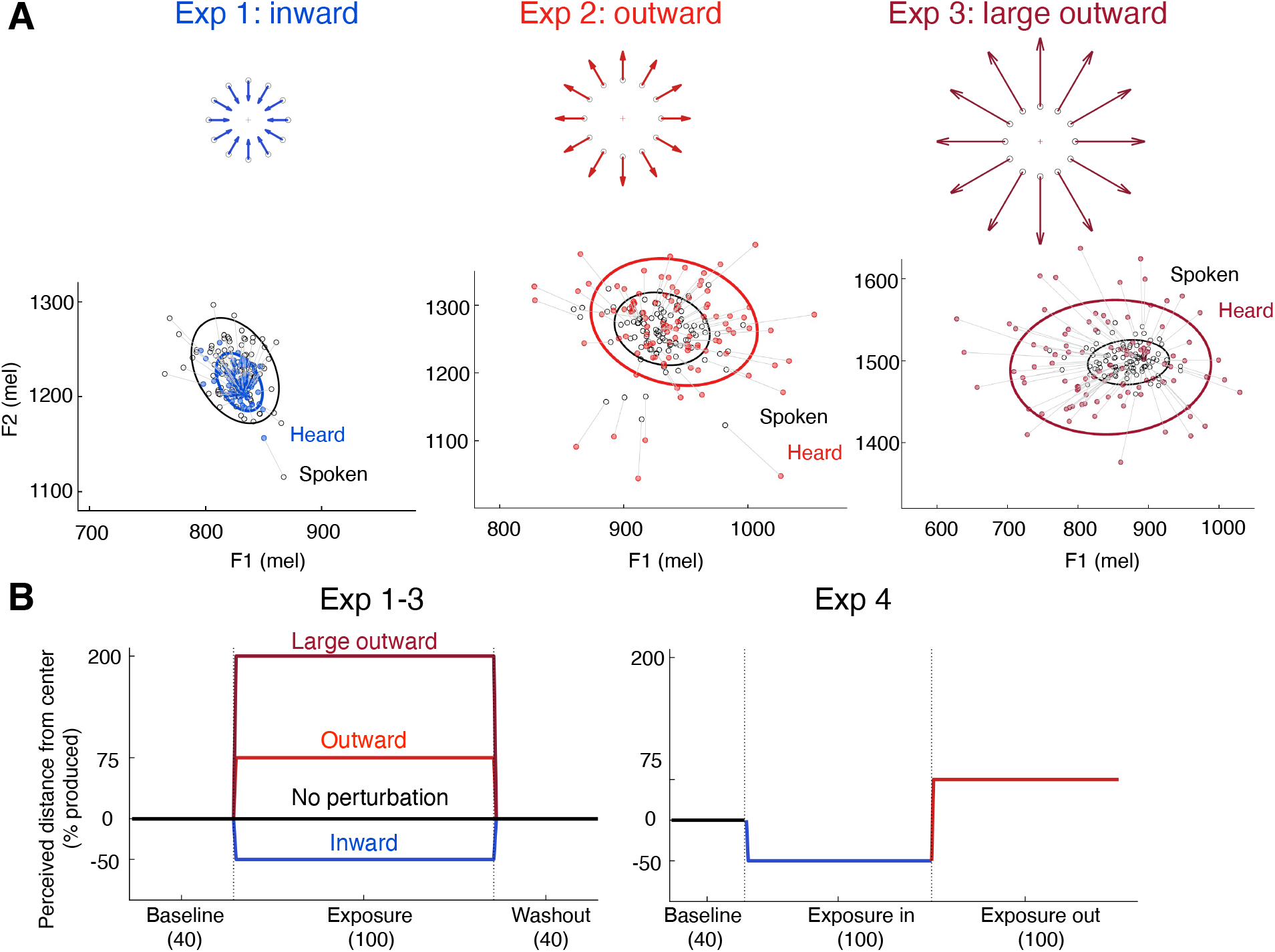
Experiment design. (A) Schematic (top) and examples from representative participants (bottom) of perturbations applied to speech vowel formants: inward-pushing (Exp 1, left), outward-pushing (Exp 2, middle), and large outward-pushing (Exp 3, right). Note different axis scales in the example data across experiments. Black circles represent the formant values participants produced and colored circles indicate what was played back to them over headphones. The ellipses represent a 95% confidence interval around the data points of the same color. (B) Experimental procedure and magnitudes of the perturbation applied in Experiments 1-3 (left panel) and Experiment 4 (right panel).

By altering participants’ perceived trial-to-trial variability without affecting their overall mean behavior, we can test whether variability in speech production is actively monitored and regulated. In a series of four experiments, we assess the effects of manipulations that both increase and decrease the perceived variability of participant’s speech behavior. We predicted that a perturbation that reduces perceived variability would lead to increases in produced variability, as participants would be free to be less precise in their production without negatively affecting their perceived accuracy. Conversely, we expected that a perturbation that increases perceived variability would have the opposite effect (i.e. lead to decreases in produced variability). Surprisingly, we found that both types of perturbation caused participants to increase their produced variability, while only the perturbation which increases perceived variability affected a behavioral measure of error sensitivity. Simulations of the applied perturbations using a well-established state space model of sensorimotor learning suggest that the increases in produced variability in response to the two types of perturbations may arise through distinct mechanisms: an increase in controlled variability in response to the perturbation that reduces perceived variability, and an increase in sensitivity to auditory errors in response to the perturbation that increases perceived variability. All together, these results suggest that motor variability is actively monitored and regulated even in complex and well-practiced behaviors, such as speech.

## Methods

### Participants

87 native speakers of American English between the ages of 18 and 66 years, with no reported history of hearing loss or neurological disorders, took part in the study (Experiment 1: N = 24, 28.8 ± 12.3 years, 18/6 females/males; Experiment 2: N = 22, 25.9 ± 9.7 years, 14/8 females/males; Experiment 3: N = 21, 33.3 ± 13.6 years, 12/9 females/males; Experiment 4: N = 20, 28 ± 12.9 years, 15/5 females/males). Participants provided informed consent and were compensated either monetarily or with course credit for their participation. The Institutional Review Board of the University of Wisconsin–Madison approved the experimental protocol.

### Apparatus

Participants were exposed to a real-time perturbation of the first and second vowel formants (F1/F2) designed to affect the perceived variability of their speech production. A modified version of Audapter (Cai et al., 2008; Tourville et al., 2013) was used to record participants’ speech, alter the speech signal when necessary, and play the (potentially altered) signal back to participants. The experiment was conducted in a quiet room with participants seated in front of a computer screen. In each trial, one of the three stimulus words (“bead”, “bad”, and “bod” in Experiment 1, 2, 4 or “bead”, “bad”, and “bed” in Experiment 3) was pseudorandomly selected and displayed on the screen, and participants read it aloud. Speech was recorded at 16 kHz via either a head-mounted microphone (AKG C520, Exp 1-2) or a desktop microphone (Sennheiser MKE 600, Exp 3-4). The output of Audapter was played back to participants via closed-back circumaural headphones (Beyerdynamic DT 770) with an unnoticeable delay of ~18 ms, as measured on our system following Kim & Max (2020). All trials were processed through Audapter in the same manner, regardless of whether a perturbation was applied. Speech was played back at a volume of approximately 80 dB SPL and mixed with speech-shaped noise at approximately 60 dB SPL, which served to mask potential perception of the participants’ own unaltered speech through air and bone conduction. The volume of speech playback varied dynamically with the amplitude of participants’ produced speech.

### Experiment-specific auditory perturbation

We designed a modified version of Audapter (Parrell & Niziolek, 2021) that is able to affect the perceived variability of speech production by specifying formant perturbations as a function of the current values of F1 and F2 (Figure 1A). A participant-specific perturbation field was calculated for each vowel such that every production during the exposure phase was shifted toward (inward-pushing perturbation) or away from (outward-pushing perturbation) the mean F1/F2 values of that participant’s distribution for that vowel (the vowel “targets”). The magnitude of the perturbation was defined as a percentage of the distance between the currently produced vowel formants and the vowel targets. By scaling the error between the vowel formants and their targets, the inward- and outward-pushing perturbations respectively reduce and magnify the perceived variability of speech production.

In Experiment 1 (**inward-pushing perturbation**), participants received a perturbation that shifted every production *towards* the vowel target. The perturbation was 50% of the distance, in F1/F2 space, between the current formant values and the vowel targets (see Figure S1).

In Experiment 2 (**outward-pushing perturbation**), participants received the opposite perturbation, a shift of every production *away* from these targets, with a slightly larger perturbation magnitude (i.e., 75% of the distance to the vowel target, Figure S1).

In Experiment 3 (**large outward-pushing perturbation**), we aimed to test the possibility that the failure to find the hypothesized reduction in variability in response to the outward-pushing perturbation applied in Experiment 2 was due to inadequate perturbation magnitude. We suspected that a larger perturbation magnitude might be needed to drive participants to produce compensatory reductions in variability. The perturbation magnitude was therefore increased to 200% of the distance to the vowel target in Experiment 3 (Figure S1) and one stimulus word was changed from a corner vowel word (“bad”) to a non-corner vowel word (“bed”) to test the effect of the perturbation on non-corner vowels.

In Experiment 4 (**inward-outward pushing perturbation**), we further examined whether limits on articulatory precision prevented participants from reducing their produced variability in response to outward-pushing perturbations in Experiment 2 and 3: that is, if individuals already produce vowels at the lower limit of variability, they may not be able to produce further variability decreases. In Experiment 4, participants experienced two exposure phases: first, to an inward-pushing perturbation, which served to increase participants’ produced variability above baseline levels, and then to an outward-pushing perturbation. The perturbation magnitudes were 50% of the distance to the vowel target for both inward-pushing and outward-pushing phases.

In order to control for potential changes in variability over the course of ~500 trials of single word production, we additionally analyzed an existing dataset (Parrell & Niziolek, 2021) with a similar experimental structure, though with no auditory perturbation applied (auditory feedback was processed through Audapter in the same manner as the baseline phases in Experiments 1-4). This control experiment included 460 trials of single word productions (115 repetitions of each stimulus word in Experiments 1, 2 and 4 — “bead”, “bad”, and “bod”—as well as the word “booed”, which did not occur in Experiments 1-4 and was not analyzed). To match the experimental design of the current study, the 460 trials were divided into four phases: baseline (30 trials per stimulus), early exposure (30 trials per stimulus), late exposure (30 trials per stimulus) and washout (25 trials per stimulus). The experimental setup for the control dataset, including recording, processing, and headphone presentation of speech, was identical to the Experiments 1-4.

### Procedure

In all experiments, stimulus words were presented on the computer screen for 1.5 s, one at a time. The interstimulus interval was randomly jittered between 0.25-1 s. Participants were instructed to read each word out loud as it appeared.

Each experiment had three phases (Figure 1B). Experiments 1-3 were divided into baseline, exposure and washout phases. In the baseline phase (40 trials per stimulus), participants received unaltered auditory feedback and we measured participants’ mean F1/F2 values for each vowel. These values were subsequently used to calculate the participant-specific perturbation field (see above). The exposure phase followed the baseline phase. In the exposure phase, participants produced each stimulus 100 times while receiving either the inward-pushing, outward-pushing, or large outward-pushing perturbation (see experiment-specific auditory perturbation above). Experiments 1-3 ended with a washout phase where participants produced each word 40 times with unaltered auditory feedback. In Experiment 4, following the baseline phase, participants experienced two sequential 300-trial (each stimulus 100 times) exposure phases, first with an inward-pushing perturbation and then with an outward-pushing perturbation. A short self-timed break was given every 30 trials in all experiments.

After they completed the experiment, participants in all four experiments were given a brief questionnaire to assess their awareness of the perturbation as well as whether they adopted any strategy and, if so, what that strategy was.

### Quantification and statistical analysis

F1 and F2 were tracked offline using wave_viewer (Niziolek & Houde, 2015), which provides a MATLAB GUI interface to formant tracking using Praat (Boersma & Weenink, 2019). Linear predictive coding (LPC) order and pre-emphasis values were adjusted individually for each participant. All trials were first checked manually for errors in production (e.g., if the participant said the wrong word). Vowel onset and offset were detected automatically using a participant-specific amplitude threshold and errors in the location of these automatically-defined landmarks were manually corrected using the audio waveform and spectrogram. Vowel onset was marked at the point where periodicity was visible in the waveform and formants were visible in the spectrogram. Vowel offset was marked at the point where formants, particularly F2 and higher, were no longer visible. Errors in formant tracking were manually corrected by adjusting the LPC order or pre-emphasis value on a trial-specific basis. In total, 1.9% of the data were excluded due to production errors or unresolvable errors in formant tracking. For each trial, F1 and F2 values were calculated by averaging formants from a 50-ms segment at both the beginning (vowel onset) and the middle (vowel midpoint) of each vowel.

The primary goal of the analysis was to test how variability changed across the different phases of each experiment. For offline analysis, the exposure phase was equally divided into early exposure and late exposure phases to make sure each phase contained a similar number of trials. Variability within each experimental phase was calculated as the average of the 2D distances in F1/F2 space between each production of a vowel and the center of the distribution for that vowel in that phase, measured from the first 50 ms of vowel. In order to test how variability may change in specific dimensions, we additionally calculated formant variability separately along the F1 and F2 axes, as well as along the major and minor axes of produced variability in the baseline phase. The variability along the F1 and F2 axes was defined as the standard deviation (SD) of F1 and F2 values of all productions of a stimulus word during each phase. For the major and minor axes of variability, an ellipse which represents a 95% confidence interval of trials in F1-F2 spaces was fitted for each stimulus word and each experimental phase using the Principal Components method. The vector representing the F1 and F2 values for each trial was projected into a component along the major axis of the fitted ellipse and a component along the minor axis perpendicular to the major axis. The variability along the major and minor axes was defined as the standard deviation (SD) of projected values along that axis of all productions of a stimulus word during each phase.

We additionally measured vowel centering, a measure of within-trial correction for variability, calculated by the change in variability from vowel onset (first 50 ms) to vowel midpoint (middle 50 ms). Vowel centering allows us to determine whether participants altered their within-trial control of variability in response to the perturbation (Niziolek & Guenther, 2013; Niziolek & Kiran, 2018; Niziolek & Parrell, 2021). Centering was measured separately for each vowel in each experimental phase.

Repeated measure analyses of variance (ANOVAs) were conducted separately for the variability and centering results and for each experiment, with phase and vowel identity as within-subject factors. For the control data and Experiments 1-3, data from baseline, late exposure and washout phases were included in the repeated ANOVAs, while data from baseline, inward-pushing exposure, and outward-pushing exposure phases were included in Experiment 4. Post-hoc comparisons (paired t-tests) were only conducted in the event of a significant main effect of phase or interaction. As an exploratory analysis, multiple regression was conducted in each experiment to determine whether produced variability changes could be predicted by baseline variability and vowel identity. Finally, variability changes along F1/F2 or major/minor axes were examined separately by three-way repeated measures ANOVAs which included phase, vowel identity and axis (i.e. F1/F2 or major/minor) as within-subject factors. The significance level for all statistical tests was p < .05, with a Bonferroni correction for multiple comparisons for post-hoc tests.

All statistical analyses were conducted in R (R Core Team, 2019). Repeated measures ANOVAs and pairwise paired t-tests were conducted with the rstatix package (Kassambara, 2021), in which partial eta squared (η_p_^2^) and Cohen’s *d* were calculated for repeated ANOVAs and paired t-tests, respectively, to determine effect size for statistically significant effects. Greenhouse-Geisser Correction was applied automatically to correct the degrees of freedom when sphericity was violated. Multiple regression models were constructed using the package stats. Data and associated code is available at https://osf.io/stjc9/. Some functions rely on additional code available at https://github.com/carrien/free-speech.

### Model simulations

In order to assess the potential mechanisms underlying the patterns of variability changes observed in Experiments 1-4, we conducted a simulation of speech behavior using a version of the well-established state space models that have been used in studies of sensorimotor adaptation to sensory perturbations in both limb (Baddeley et al., 2003; Donchin et al., 2003; Thoroughman & Shadmehr, 2000) and speech (Daliri, 2021; Daliri & Dittman, 2019) motor control. Importantly, analogous models have also been used successfully in studies of sensorimotor corrections for self-produced variability in reaching (Ahn et al., 2016; Blustein et al., 2021; Scheidt et al., 2001), similar to the current experiments. The model is:

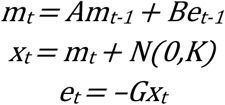

This model assumes that the intended production (*m*) on a given trial (*t*) aims to achieve a particular target (by convention, set to 0) based on a weighted contribution of the production on the previous trial (*t* – 1) and the error (*e*) experienced on the previous trial. *A* is the forgetting factor that determines the contribution of the previous trial, and *B* represents the sensitivity of the system to errors. The error *e* is the difference between the actual production outcome (%) and the target, where the production *x* is the result of the intended production *m* plus noise (*N*) drawn from a gaussian distribution with a mean of 0 and a standard deviation of *K*. In order to account for the gains applied to the auditory feedback in this experiment, the observed outcome *x* is multiplied by a gain factor *G* to derive the error. Aside from the addition of the gain factor *G*, this is identical to the formulation in Ahn et al., (2016), if *A* = 1.

While the state space model is often used to model sensorimotor adaptation to external perturbations, the addition of the noise term on the final motor output also permits modelling of correction for self-produced variability (Ahn et al., 2016; Blustein et al., 2021; Scheidt et al., 2001). For simplicity, we initially set *A* to 1 (Ahn et al., 2016) and *B* to 0.1 given previous results fitting state space models to adaptation in speech (Daliri, 2021; Daliri & Dittman, 2019). We then estimated an initial value of *K* that would generate an observed distribution similar to the experimentally observed variability in the baseline phase across experiments (roughly 30 mels). Because the errors in this model are generated stochastically, we need a large number of simulations to achieve an accurate estimate of the underlying distribution of variability that would be generated through this model. To that end, we ran batches of 1000 simulations while varying *K*. Each simulation consisted of 40 trials, equivalent to the number of trials used to estimate variability in our experimental data. Our initial simulations indicated that setting *K* to 30 resulted in a mean observed variability close to the observed values in the baseline phase across experiments. Varying *A* within the range reported in (Daliri, 2021) had very minor effects on the results; the effects of varying *B* are explored below.

Our goal in modeling was to assess the potential causes of the increase in variability observed in the behavioral data in Experiments 1-4. To do this, we systematically varied the underlying variability *K* as well as the sensitivity to errors *B. K* varied from 30 to 40 in steps of 0.5. *B* varied from 0 to 1 in steps of 0.05, where 0 would represent no correction for observed errors and 1 would represent full correction. For each step of *K* and *B*, 1000 simulations (40 trials each) were conducted. The standard deviation of *x* was calculated for each simulation, and the mean of these standard deviations was calculated to generate an estimate of the expected variability with that particular parameter set. We ran separate simulations for the inward-pushing perturbation in Experiment 1 (setting *G* to 0.5) and the outward-pushing perturbation in Experiment 2 (setting *G* to 1.75).

## Results

### Overall variability changes

Over four experiments, we implemented an auditory perturbation designed to increase or decrease participants’ perceived trial-to-trial variability without affecting their overall mean. To confirm that this was achieved, the perturbations applied to F1 or F2 frequencies during the exposure phase in each experiment were averaged for each participant and compared against zero (no mean formant value change) using one-sample t-tests. The mean F1 perturbations ranged between 2.5 and 13.4 mels in Experiments 1-4 and significantly differed from zero only in Experiment 3 (large outward-pushing, mean F1 perturbation: 13.4 mels, *t*(20) = 2.47, *p* = 0.023). Similarly, the mean F2 perturbations ranged between −1.23 and 12.39 mels in Experiments 1-4, with no significant difference from zero in any case (uncorrected p > 0.05). It is worth mentioning that the just noticeable difference (JND) in F1 and F2 for isolated English vowels is around 14 mels and 20 mels, respectively (Kewley-Port & Watson, 1994).

We then measured how participants changed their produced variability after the perceived variability had been increased or decreased. As a control, we analyzed an existing dataset (Parrell & Niziolek, 2021) with no auditory perturbation. As expected, the control group showed no change in variability over the course of the experiment (Figure 2A, *F*(2,48) = 0.434, *p* = 0.650).

**Figure 2.**
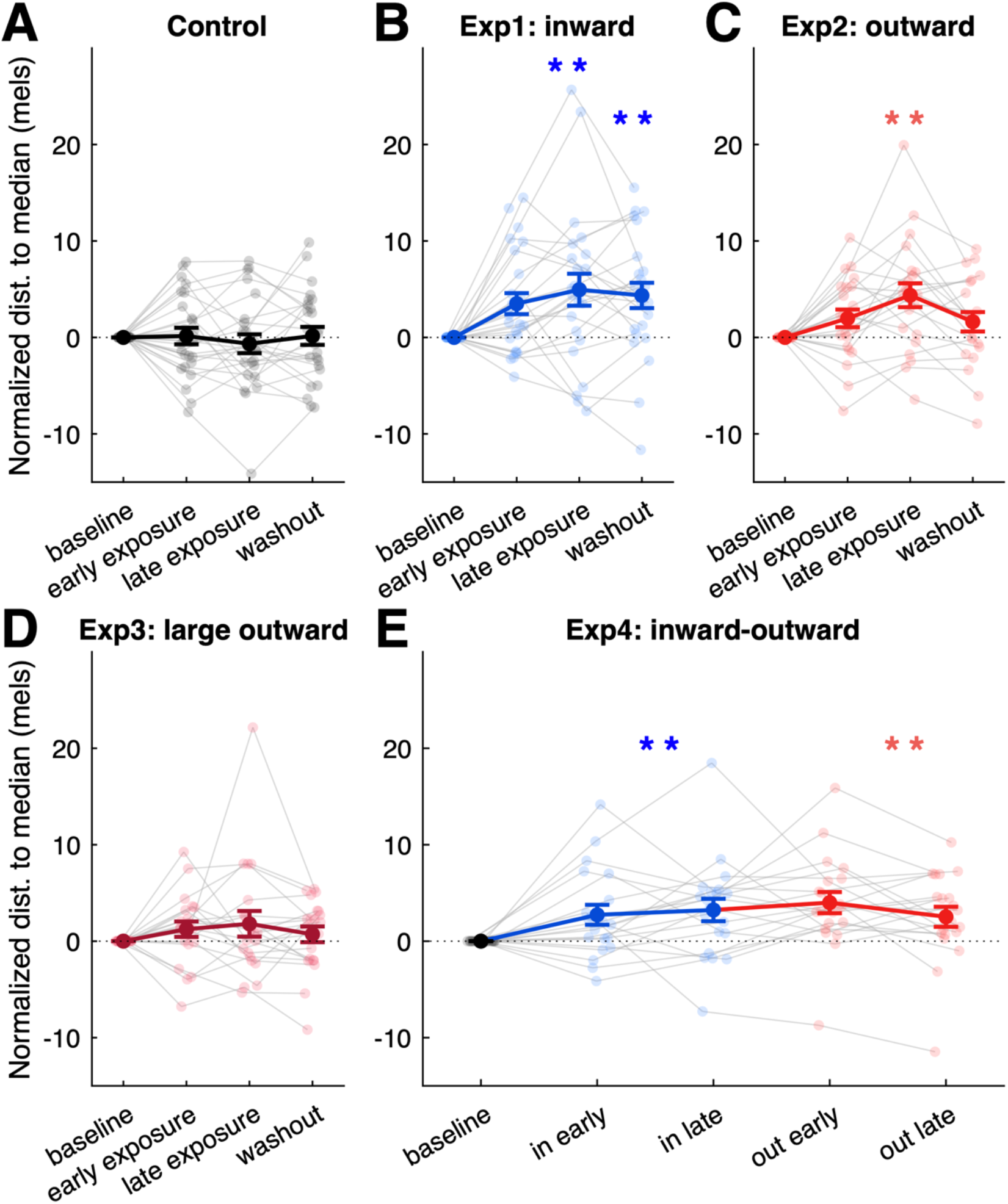
Baseline-normalized variability changes. Individual (small transparent dots, thin lines) and group means (large solid dots, thick lines) of baseline-normalized variability (normalized by subtracting the average value in the baseline from the remaining trials) in the baseline, exposure and washout phases. Error bars show standard error. ** indicates significant change (p < 0.001) from baseline.

Participants exposed to the **inward-pushing perturbation (Experiment 1)** increased produced variability (Figure 2B, main effect of phase: *F*(2,46) = 7.55, *p* < 0.001, *η_p_*^2^ = 0.25) while the perturbation was applied (+5.0 mels, *t*(71) = 4.59, *p* < 0.001, *d* = 0.54) as well as after it was removed (+4.4 mels, *t*(71) = 4.89, *p* < 0.001, *d* = 0.58). Unexpectedly, participants exposed to the **outward-pushing perturbation (Experiment 2)** also increased produced variability (Figure 2C, main effect of phase: *F*(2,40) = 7.93, *p* < 0.001, *η_p_*^2^= 0.28). However, unlike the maintained variability change seen in the inward-pushing perturbation, the increased variability during exposure (+4.3 mels, *t*(65) = 4.10, *p* < 0.001, *d* = 0.50) returned to near-baseline levels when the outward-pushing perturbation was removed (+1.6 mels, *t*(62) = 1.93, *p* = 0.175).

We reasoned that the failure to find the hypothesized reduction in variability in response to the outward-pushing field could potentially be attributed to the size of the perturbation used. Although this perturbation was slightly larger than in Experiment 1 (75% vs 50% of the distance to vowel target), it may still have been too small to induce participants to reduce their variability. **Experiment 3 (large outward-pushing perturbation)** aimed to delineate this by testing another group of participants who received outward-pushing perturbations with a 200% increase in distance to the center of vowel distribution (vs 75% in Experiment 2). However, participants exposed to this large outward-pushing perturbation did not change their produced variability (Figure 2D, main effect of phase: *F*(1.55,30.95) = 1.39, *p* = 0.26).

Together, the results of both experiments employing outward-pushing perturbation fields (Experiment 2 and 3) suggest these perturbations do not drive participants to produce compensatory reductions in variability. One possibility for this behavior is that speech movements are already produced at or near the lower limit of an individual speaker’s precision ability. **Experiment 4 (inward-outward pushing perturbation)** aimed to test this possibility by examining whether an outward-pushing perturbation can reduce produced variability of participants *after* their variability has increased above baseline levels due to exposure to an inward-pushing perturbation as seen in Experiment 1. Participants significantly changed their produced variability during the course of Experiment 4, reflected by a main effect of phase (Figure 2E, *F*(2,38) = 9.71, *p* < 0.001, *η_p_*^2^= 0.34). As expected given the results of Experiment 1, variability increased when the inward-pushing perturbation was applied (+4.1 mels, *t*(59) = 4.77, *p* < 0.001, *d* = 0.62). However, participants did not change their produced variability back to the baseline when receiving the outward-pushing perturbation in the following phase (+4.1 mels, *t*(59) = 4.13, *p* < 0.001, *d* = 0.53). While this result replicates the increase in variability observed during the exposure phase in Experiments 1 and 2, it suggests the failure to find the expected reduction in variability in Experiments 2 and 3 was not caused solely by a “lower limit” on variability.

Finally, while not the primary focus of the study, we evaluated whether participants adjusted their variability differently along the major/minor axes of variation using three-way repeated ANOVAs including three within-subject factors (phase, vowel, axis). This analysis replicated the results of overall variability changes (distances in F1-F2 spaces): a significant main effect of phase was found in Experiments 1, 2, 4, but not in the control data or in Experiment 3. Perhaps more interestingly, no significant two-way interaction between phase and measure was found in any of these experiments, suggesting participants adjusted their variability along the major- and minor-axis similarly. Similar results were found from models of F1 and F2 variability. See Supplemental Material, Figures S2, S3 and Tables S1, S2 for detailed results and statistics.

### Centering changes

In order to determine whether participants adjusted their within-trial control of variability in response to the perturbation, we additionally measured vowel centering (Niziolek & Guenther, 2013; Niziolek & Kiran, 2018; Niziolek & Parrell, 2021), the reduction in variability from vowel onset (first 50 ms) to vowel midpoint (middle 50 ms). Similar to the analyses on variability, the control group did not show any change in centering over the course of the experiment (Figure 3A, *F*(2,48) = 1.45, *p* = 0.244). However, unlike the overall variability changes observed above, no change in centering was seen in participants who received the **inward-pushing perturbation** (Experiment 1, Figure 3B, *F*(1.49, 34.27) = 0.06, *p* = 0.896). In contrast, centering did increase when participants were exposed to the **outward-pushing perturbation** (Experiment 2, Figure 3C, main effect of phase: *F*(2,40) = 3.94, *p* = 0.027, *η_p_*^2^= 0.165), suggesting these participants became more responsive to errors. This increase in centering during exposure (+ 2.2 mels, *t*(65) = 3.33, *p* = 0.004, *d* = 0.41) was not retained during the washout phase (+ 0.1 mels, *t*(62) = 0.11, *p* = 1.000). Contrary to the increase in centering observed in Experiment 2, participants who received a **large outward-pushing perturbation** in Experiment 3 (i.e. 200% increase in distance to the vowel target) did not show an increase in centering over the course of experiment (Figure 3D, *F*(2,40) = 0.87, *p* = 0.428), actually tending to slightly decrease centering during and after exposure. In Experiment 4 **(inward-outward pushing perturbation)**, participants showed changes in centering over the course of the experiment, indicated by a marginally significant main effect of phase (Figure 3E, *F*(1.4, 26.5) = 3.15, *p* = 0.075, *η_p_*^2^= 0.142). More specifically, the results replicated the patterns observed in Experiments 1 and 2: the initial inward-pushing perturbation did not induce any significant increase in centering (+ 0.53 mels, *t*(59) = 0.38, *p* = 1. 000), while the subsequent outward-pushing perturbation did (+ 2.7 mels, *t*(59) = 2.27, *p* = 0.05, *d* = 0.29).

**Figure 3.**
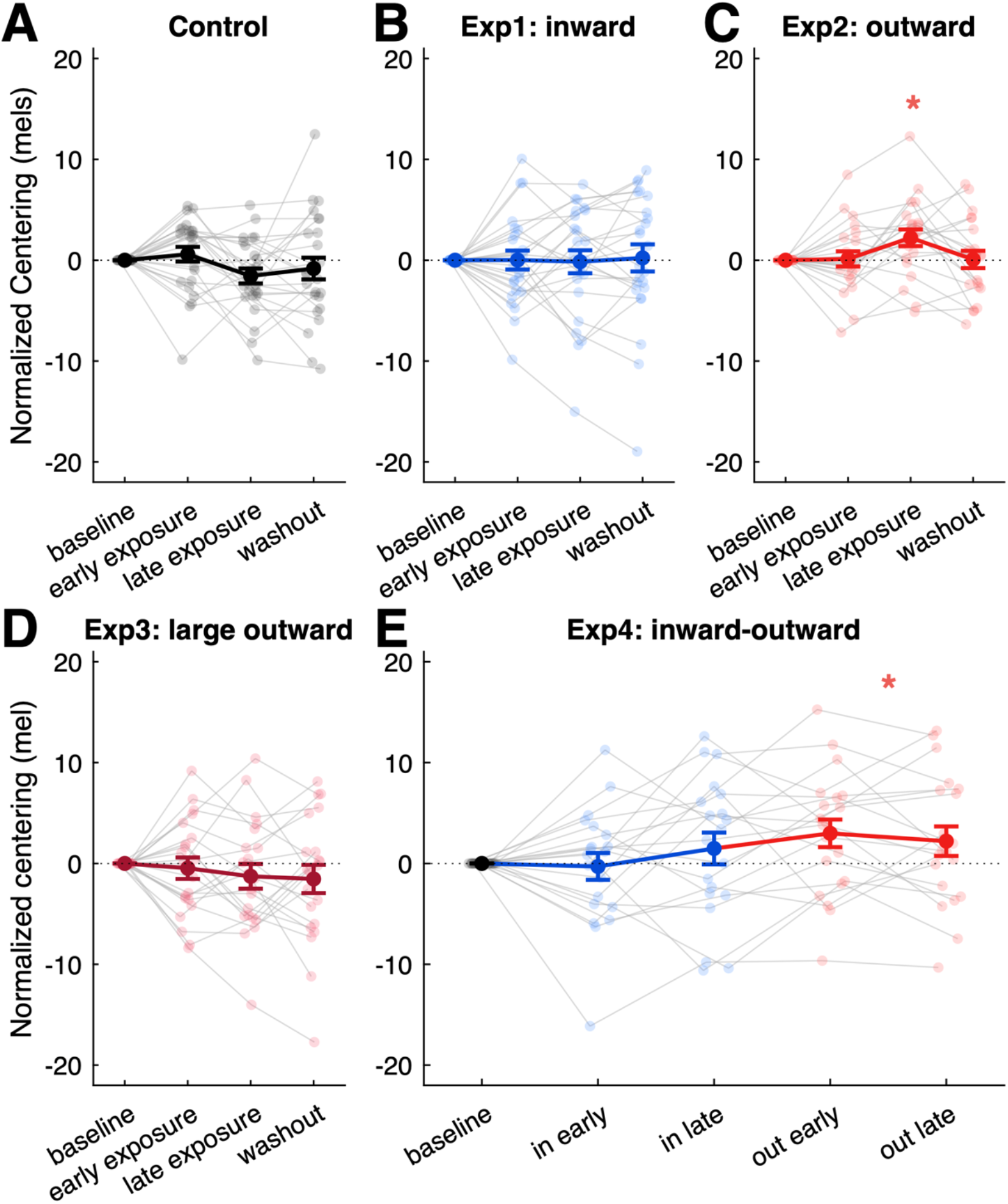
Baseline-normalized centering changes. Individual (small transparent dots, thin lines) and group means (large solid dots, thick lines) of baseline-normalized centering (normalized by subtracting the average value in the baseline from the remaining trials) in the baseline, exposure and washout phases. Error bars show standard error. * indicates significant change (p < 0.05) from baseline.

### Correlation between variability change and baseline variability

Multiple regression was conducted in each experiment to determine whether changes in produced variability could be predicted by baseline variability and vowel identity at the individual level (see Figure 4). Results showed that baseline variability was not predictive of the individual change in variability observed during the late exposure phase in either the control (*β* = −0.18, *p* = 0.094) or Experiment 2 (**outward-pushing perturbation**, *β* = −0.024, *p* = 0.846). However, this relationship was observed in the Experiment 1 (**inward-pushing perturbation**, *β* = −0.44, *p* < 0.001), such that participants with lower variability in the baseline phase showed larger variability increases. In Experiment 3 **(large outward-pushing perturbation)**, where there was no consistent change in variability over the course of the experiment, baseline variability was nonetheless predictive of individual changes in variability (*β* = −0.38, *p* = 0.002): participants with higher variability in the baseline phase tended to decrease variability, and vice versa. Experiment 4 (**inward-outward pushing perturbation**) replicated the result from Experiment 1: a significant correlation was observed between baseline variability and variability change induced by inward-pushing perturbation (*β* = −0.19, *p* = 0.049). Perhaps surprisingly given the results of Experiment 2, a similar relationship between baseline variability and variability change was also observed during later outward-pushing perturbation (*β* = −0.37, *p* < 0.001). However, it should be noted that there was a highly significant correlation in variability between the two perturbation phases (*β* = 0.54, *p* < 0.001), suggesting that the correlation observed during the outward-pushing perturbation is likely a carry-over effect of the inward-pushing perturbation which happened between baseline and outward-pushing phases.

**Figure 4.**
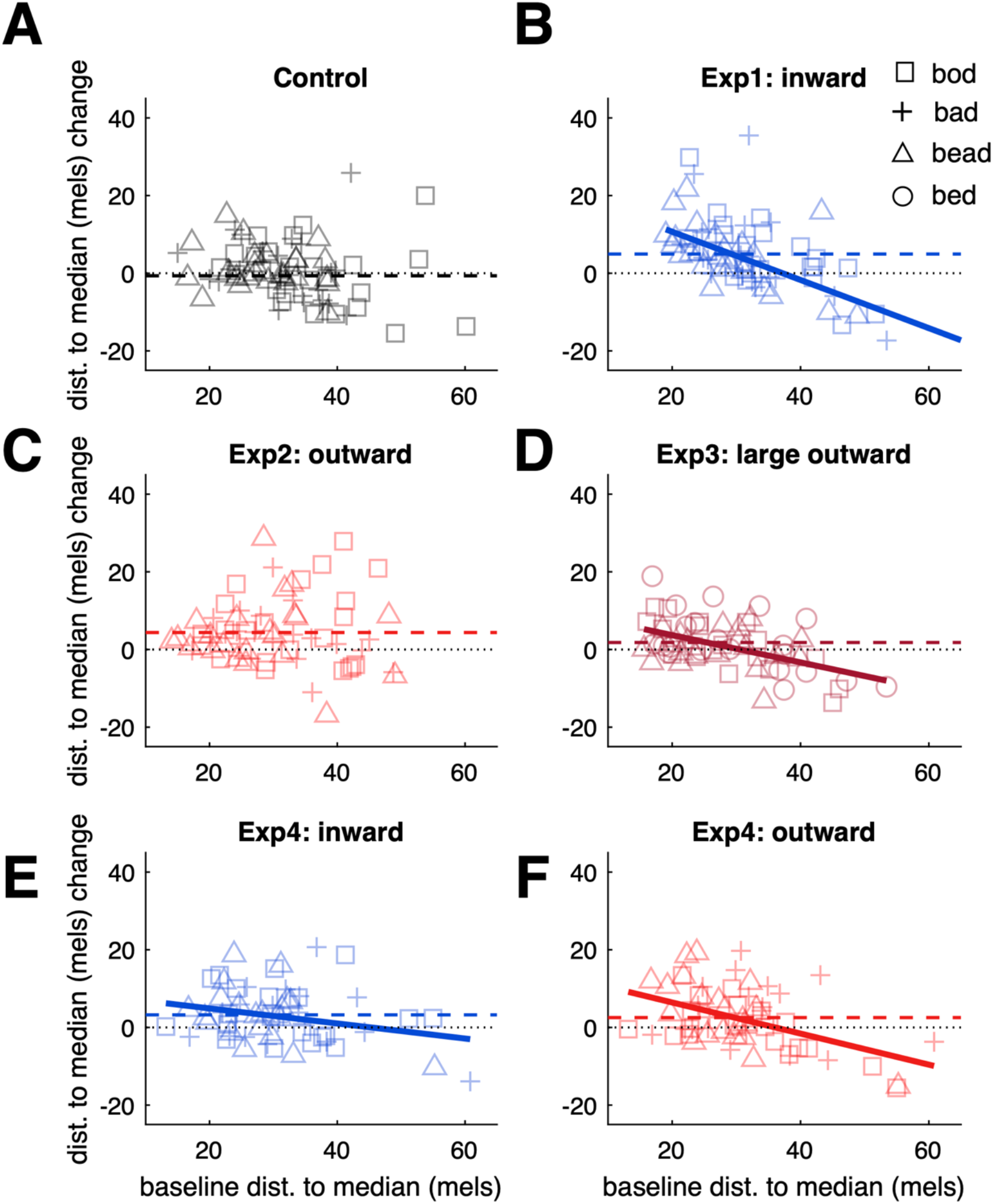
Correlation between average baseline variability and variability change across individuals. Each data point represents the average production of one stimulus word, indicated by different markers. Each participant thus contributed three data points. Note that the stimulus words were “bead”, “bad”, and “bed” in Experiment 3, and “bead”, “bad”, and “bod” in the other experiments. Significant correlations are indicated by solid lines showing the least-squares fit to the data points. The group-averaged (baseline-normalized) variability changes are indicated by colored dashed lines.

### State space simulations

Using a modified version of the typical state space model that has been shown to account for observed motor variability in reaching, we systematically varied the magnitude of motor variability (*K*) and the sensitivity of the system to observed errors (*B*). Results of these simulations are shown in Figure 5. We found that, predictably, increases in the underlying motor variability increased the observed variability in motor production for both the inward- and outward-pushing perturbations (Figure 5C). Values of *K* near 35 produced observed variability measures consistent with those in both Experiments 1 and 2 (Figure 5D). Conversely, we found that changes in error sensitivity had differential effects on motor variability depending on the perturbation field. For the inward-pushing perturbation, variation in error sensitivity had very minor effects on motor output. Conversely, for the outward-pushing perturbation, increases in error sensitivity led to large increases in motor variability (Figure 5E). Values of *B* near 0.3 resulted in a good match for the observed variability in Experiment 2, while no values of *B* provided a good match for Experiment 1 (Figure 5F). In the latter case, the maximum increase in observed variability over baseline was 3.7 where *B* = 1, still substantially less than the experimentally observed value. These results indicate that the observed increase in Experiment 1 can only be explained by an increase in the underlying motor variability, while the increase observed in Experiment 2 could arise through either an increase in controlled variability or an increase in the sensitivity to auditory errors.

**Figure 5.**
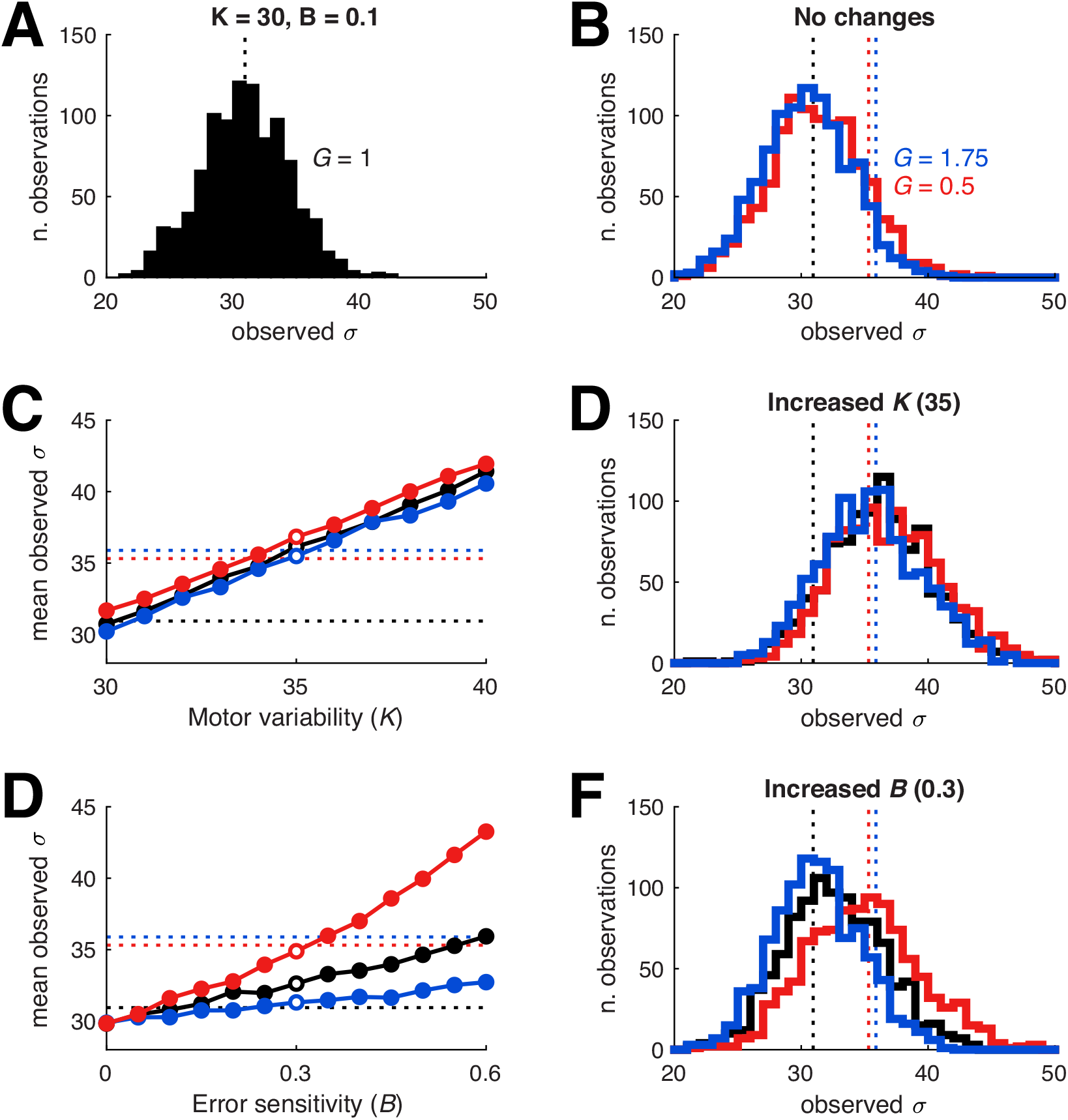
State space simulations of motor variability. For all panels, the dashed black line represents the mean variability with model parameters set to match participant behavior in the baseline phase, the dashed blue line represents the increase in variability observed in Experiment 1 (4.9 mels above baseline), and the dashed red line represents the increase in variability observed in Experiment 2 (4.4 mels above baseline). (A) shows distributions of observed variability for 1000 simulations where the gain applied to the observed error is 1, simulating the baseline phase in both Experiments 1 and 2. (B) shows similar distributions from simulations with gains set to 0.5 (blue) and 1.75 (red), mirroring the perturbations applied in Experiment 1 and Experiment 2, respectively, with no changes to other model parameters. (C) shows the result of systematically varying underlying motor variability K on observed variability with gains set to 1(black), 0.5 (blue), and 1.75 (red). The open circles in the left panel represent the value for K used in the simulations shown in (D). (D) shows the distribution of observed variability with K set to 35, chosen to roughly match the increase in variability observed in Experiments 1 and 2. (E) as for (C), showing the effect of varying the error sensitivity parameter B. Values of B greater that 0.6 (not shown) resulted in values observed variability >50 where G = 1.7 and < 34 where G = 0.5. (F) shows the distribution observed with B set to 0.3, chosen to match the increase in variability observed in Experiment 2.

### Awareness of perturbation

Most of the participants in Experiment 1, 2, and 4 (41/66) were not aware of the perturbation applied to their vowels. Although 25/66 did indicate that they thought their speech was somehow manipulated, only 1 participant correctly identified it as a change to their vowels/consonants. In contrast, nearly all participants (19/21) in Experiment 3 (large outward-pushing) reported that they thought they received a perturbation, and almost half (9/21) correctly identified it as a manipulation to their vowels. No participants reported a strategy that aimed to address the applied perturbation in any of the four experiments.

## Discussion

In a set of four experiments, we examined whether variability can be actively regulated in a complex, well-practiced motor task: speech production. Specifically, we examined whether participants would adjust their produced variability when they were exposed to real-time auditory perturbations that increased or decreased their perceived variability. Our results showed that introducing a perturbation that reduces perceived variability (Experiment 1: inward-pushing) leads to increases in produced vowel variability that remain even when normal feedback is restored, suggesting that variability is monitored and regulated over relatively long time scales. Perhaps surprisingly, a perturbation that increases perceived variability (Experiment 2: outward-pushing) also increased produced variability, though such variability change was not maintained when the perturbation was removed. In Experiment 3, participants who received a large outward-pushing perturbation (i.e. a 200% increase in distance to the vowel target) did not change their produced variability over the course of the experiment. This rules out the possibility that the failure to find the hypothesized reduction in variability in response to the outward-pushing perturbation (Experiment 2) is due to inadequate perturbation magnitude. Finally, we found that an outward-pushing perturbation cannot reduce participants’ produced variability even *after* their variability has increased above baseline levels as a result of the inward-pushing perturbation (Experiment 4: inward-outward pushing).

To our knowledge, this is the first study showing that variability in speech production can be actively controlled, consistent with recent theories that have highlighted the adaptive value of motor variability in other motor domains (Davids et al., 2003; Herzfeld & Shadmehr, 2014; Tumer & Brainard, 2007). From this perspective, motor variability can be actively generated, regulated and used by the brain to improve motor performance, reduce costs and explore new solutions (Shadmehr et al., 2016). Early evidence shows that skilled performers are able to upregulate the level of motor variability in each joint of the upper arm to meet the change of task demands/constraints, while less skilled performers, in comparison, tend to have rigidly fixed motor variability that is not fine-tuned with task constraints (Arutyunyan et al., 1968, see Newell & Vaillancourt, 2001 for a review). More recent work using computational models also found that force variability and the resulting kinematic variability are not generated primarily by random “motor noise”, and emphasize the importance of other sources of force variability which can be tuned as needed by distributed sensorimotor systems (Nagamori et al., 2021). The results from the current study extend previous work and provide support for this perspective by showing the motor system closely monitors sensory variability and uses such information to actively regulate the motor variability, even in complex and well-practiced behaviors such as natural speech.

Surprisingly, participants exposed to both inward-pushing and outward-pushing perturbations increased their produced variability. These results could be interpreted as a general variability increase induced by either repetitive productions of many utterances or a non-specific auditory perturbation (that is, any kind of auditory perturbation would lead to an increase in formant variability). However, the results from our data as well as previous studies indicate that both of the possibilities are unlikely. First, in analyzing an existing dataset where participants produced 460 utterances with normal auditory feedback (Parrell & Niziolek, 2021), we found that speakers did not significantly change their produced variability. This confirms that under normal circumstances speakers produce vowels with a level of variability that is relatively stable over time. Second, previous work has shown that participants do not change their formant variability in response to consistent auditory perturbation of both F1 and F2 (i.e. shifts of 240 Hz in F1 and 300 Hz in F2) (Nault & Munhall, 2020). Results from Experiment 3 are consistent with this result, as participants who received a large outward-pushing perturbation did not exhibit any significant change in their variability. These results, taken together, rule out the possibility of a general non-specific increase in variability induced by auditory perturbations.

In our experimental data, the increased variability induced by the auditory perturbation was maintained even when normal feedback was restored in Experiment 1, but not in Experiment 2. This difference suggests different mechanisms may have led to the increased variability observed in the two experiments. As an attempt to disentangle these potential mechanisms, we conducted a simulation using a state space model of error correction. Model simulations identified two distinct mechanisms that could lead to the observed increase in produced variability: an increase in controlled variability or an increase in the sensitivity to auditory errors. More specifically, these simulations indicated that the observed increase in Experiment 1 (inward-pushing) can only be explained by an increase in controlled variability, while the increase observed in Experiment 2 (outward-pushing) could arise through either of these two mechanisms. Importantly for Experiment 2, a return to unperturbed auditory feedback in the washout phase would not necessarily cause an immediate decrease in controlled variability, but would directly lead to a change in variability related to trial-to-trial correction for errors even with a constant error sensitivity (see Figure 5). These modeling results suggest the increase in observed variability in response to the inward-pushing perturbation in Experiment 1 was likely driven by a relaxation of controlled variability, as the perturbation “frees” the motor system to be more variable without any loss in perceived accuracy. Conversely, the results from the Experiment 2 are most consistent with an increase in error sensitivity caused by the outward-pushing field rather than a change in controlled variability. Consistent with this difference, we found a significant correlation between produced variability changes and baseline variability in Experiment 1 but not in Experiment 2. One possible explanation for the pattern observed in Experiment 1 is that participants who are naturally more variable may take advantage of the natural consequences of the inward-pushing perturbation to reduce overall perceived variability, while less variable individuals may relax their (presumably stricter) regulation of variability without negatively affecting their perceived variability. In contrast, in Experiment 2, we would not expect that changes in error sensitivity would be related to levels of baseline variability. In brief, the combined results of behavioral and model simulation point to the two distinct mechanisms that may have led to the observed increase in variability in response to inward-pushing and outward-pushing perturbation.

It is worth mentioning that while it would be ideal to compare the predictions generated by the state space model to empirical estimates of error sensitivity, limitations of the experimental design prevent us from being able to estimate such parameters. Even with recent advances in this regard (Blustein et al., 2021), accurate estimation of these parameters requires 100-200 sequential trials. In all experiments, participants produced 40-50 repetitions of each word per phase, in a pseudo-randomized order. Thus, our protocol provides both an insufficient number of trials as well as a potential confound of producing multiple intervening movements between repetitions of the same item. Future work should explore whether current methods that are known to provide accurate estimates of error sensitivity in reaching could be applied to speech motor control with a more appropriate experimental design.

We additionally measured vowel centering, a measure of within-trial correction for variability (Niziolek & Guenther, 2013; Niziolek & Kiran, 2018; Niziolek & Parrell, 2021). Unlike the overall variability changes observed in both kinds of perturbation, an increase in centering was only seen in participants who received the outward-pushing perturbation in Experiment 2 and in the outward-pushing phase of Experiment 4. This suggests that participants exposed to outward-pushing perturbations became more responsive to within-trial errors (i.e. larger within-trial feedback gains). Although it is not clear that such an increased sensitivity to errors for within-movement corrections is directly related to the error sensitivity related to cross-trial changes (i.e. gains on trial to trial learning), it is possible error sensitivity is shared across both processes. This would be consistent with the fact that we see increased centering only in Experiment 2, precisely where modeling results suggest an increase in trial-to-trial error sensitivity.

The only case where we did not see an increase in centering or variability was Experiment 3 (large outward-pushing perturbation). It is possible that the larger perturbations used in this experiment were discounted by the sensorimotor system, such that they had limited effects on speech production. This is consistent with previous studies that have similarly shown attenuated responses to large auditory feedback perturbations (Burnett et al., 1998; Scheerer et al., 2013), potentially because large perturbations are treated as externally induced, rather than self-produced errors (Korzyukov et al., 2017). This is also in line with the perturbation awareness results: almost half of the participants in Experiment 3 correctly recognized the perturbation applied on vowels, while only one participant did across all other experiments.

Together, the results of Experiments 2-4 suggest that a sensory perturbation that increases perceived variability does not drive participants to produce compensatory reductions in speech variability, contrary to previous results in non-speech motor control which showed task-relevant variability can be reduced when needed or after repeated practice (Kang et al., 2004; van Beers et al., 2013; Wong et al., 2009). One potential explanation for our failure to observe the hypothesized decrease in variability in response to the outward-pushing perturbation in Experiment 2 is that speech is already produced at the lower limits of possible task-related variability, consistent with the predictions of the Uncontrolled Manifold hypothesis and Optimal Feedback Control (Harris & Wolpert, 1998; Scholz & Schöner, 1999; Todorov, 2004). This could also explain the lack of change in produced variability when the outward-pushing perturbation occurred after variability had increased above baseline levels (Experiment 4): it is possible that controlled variability here returned to baseline levels (without reducing past that point), but was counteracted by the increase of error sensitivity induced by outward-pushing perturbations.

Another possibility is suggested by the fact that in reaching tasks, only task-relevant variability has been observed to reduce experimental tasks, while task-irrelevant variability remained high (Robertson & Miall, 1997; Scholz & Schöner, 1999). Thus, it is possible that participants in the current experiments reduced variability along a particular dimension, even if their overall variability increased. To explore this possibility, we calculated changes in variability along participant-specific major and minor axes of variability in the baseline phase, assuming that the minor axis of variability may be more tightly controlled (Scholz & Schöner, 1999). However, we found that participants produced similar changes in variability along the major- and minor-axis, suggesting speech motor system might control variability more globally compared to the selective regulation in certain components of movement variability observed in non-speech motor control (Abe & Sternad, 2013; Sternad et al., 2011). It is important to note, however, that the major- and minor-axis in speech production are not necessarily equivalent to the task-relevant and task-irrelevant dimensions in non-speech control, and indeed may both contribute to task performance. Further work is needed to clarify this point.

In summary, we have shown that individuals modify their produced variability when their perceived trial-to-trial variability is altered. Decreases in perceived variability lead to increases in produced variability, likely due to loosened restrictions on variability production, particularly in individuals with inherently low variability. These changes are retained even when the perturbation is removed, suggesting that the monitoring and regulation of variability acts relatively slowly. Conversely, variability also increases in response to perturbations which increase perceived variability, potentially due to increases in error sensitivity as participants try to correct for the perceived errors. Together, these results are consistent with recent evidence that suggests motor variability should be viewed as an important feature of how the sensorimotor system operates and learns rather than as the inevitable and unintended consequence of motor noise. Our results also highlight the importance of having a better understanding of motor variability during speech production, which has been largely overlooked in current theories and models of speech motor control.

## Acknowledgments

This work was supported by NIH grant R01 DC019134 and a grant awarded through the University of Wisconsin–Madison Fall Research Competition.

## Supplementary materials

**Figure S1.**
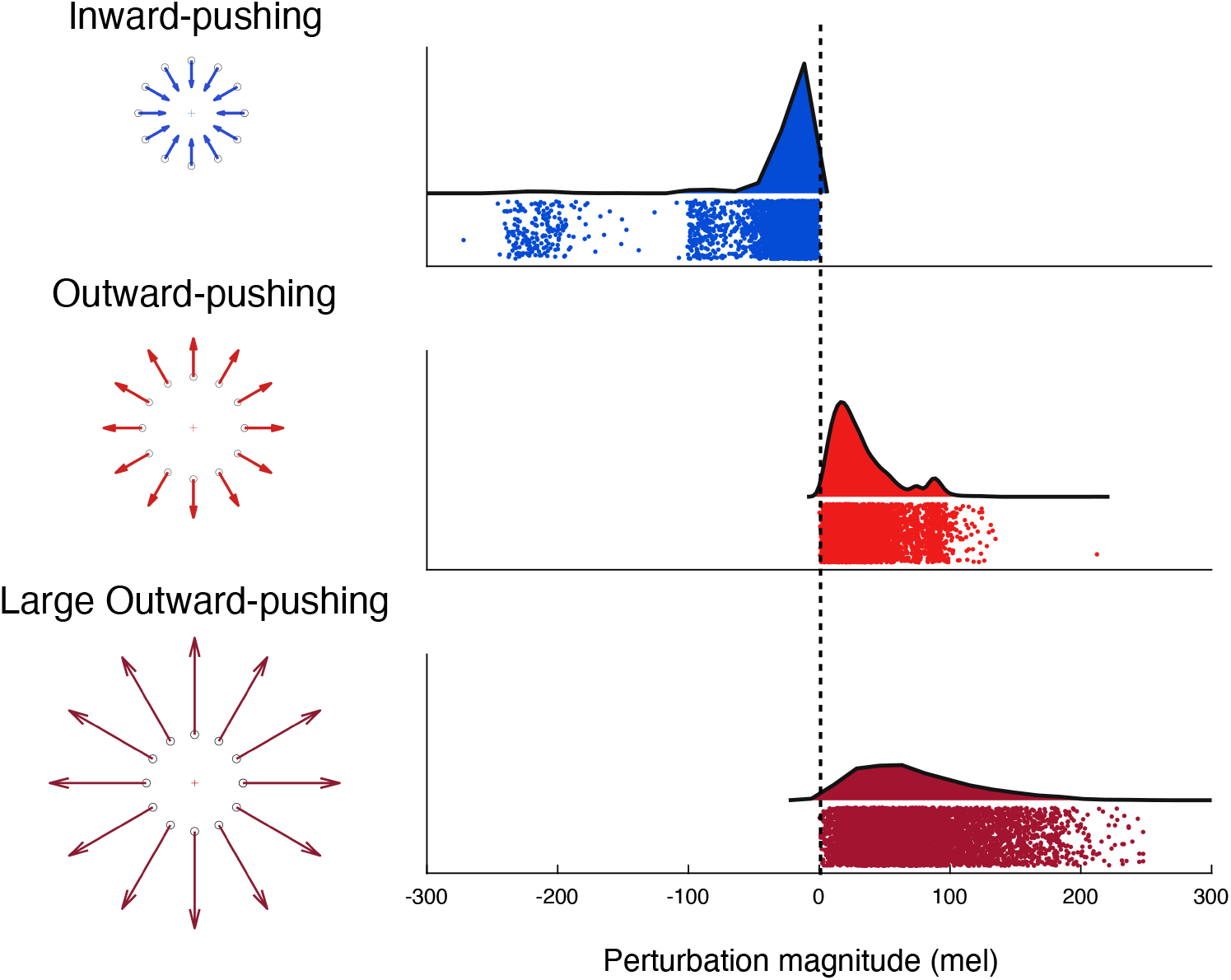
Schematic (left) and raincloud plot (right) of perturbation applied to speech vowel formants: inward-pushing pushing (Exp 1, top), outward-pushing (Exp 2, middle), and large outward-pushing (Exp 3, buttom). The raincloud plots show the magnitude of the applied perturbations both as a distribution (the ‘cloud’) and with jittered raw data (the ‘rain’).

**Table S1.** Variability (SD) changes along major and minor axis.

|  | Control | Experiment 1 | Experiment 2 | Experiment 3 | Experiment 4 |
| --- | --- | --- | --- | --- | --- |
| **Main effect:** |  |  |  |  |  |
| Phase | F(2, 48) = 0.45,<br>p = 0.642 | F(2, 46) = 9.22,<br>p < 0.001** | F(2, 40) = 5.74,<br>p = 0.006* | F(1.5, 29.9) = 1.36,<br>p = 0.267 | F(2, 38) = 15.41,<br>p < 0.001** |
| Vowel | F(2, 48) = 5.39,<br>p = 0.008* | F(2, 46) = 17.31,<br>p < 0.001** | F(2, 40) = 17.82,<br>p < 0.001** | F(1.3, 25.2) = 1.28,<br>p = 0.28 | F(2, 38) = 7.47,<br>p = 0.002* |
| Axis | F(1, 24) = 110.13,<br>p < 0.001** | F(1, 23) = 116.54,<br>p < 0.001** | F(1, 20) = 40.02,<br>p < 0.001** | F(1, 20) = 61.5,<br>p < 0.001** | F(1, 19) = 77.23,<br>p < 0.001** |
| **Two-way interaction:** |  |  |  |  |  |
| Phase × Vowel | F(4, 96) = 1.90,<br>p = 0.117 | F(4, 92) = 0.60,<br>p = 0.662 | F(4, 80) = 2.01,<br>p = 0.101 | F(2.1, 41.4) = 0.77,<br>p = 0.474 | F(4, 76) = 1.06,<br>p = 0.385 |
| Phase × Axis | F(2, 48) = 0.83,<br>p = 0.443 | F(2, 46) = 2.84,<br>p = 0.069 | F(1.6, 31.5) = 0.33,<br>p = 0.669 | F(1.5, 29.4) = 0.28,<br>p = 0.693 | F(2, 38) = 0.78,<br>p = 0.465 |
| Vowel × Axis | F(2, 48) = 5.87,<br>p = 0.005* | F(2, 46) = 4.21,<br>p = 0.021* | F(1.5, 29.2) = 0.183,<br>p = 0.185 | F(1.2, 24) = 0.05,<br>p = 0.858 | F(2, 38) = 8.93,<br>p < 0.001* |
| **Three-way interaction:** |  |  |  |  |  |
| Phase × Vowel × Axis | F(4, 96) = 1.69,<br>p = 0.160 | F(2.4, 54) = 3.07,<br>p = 0.047* | F(4, 80) = 0.58,<br>p = 0.681 | F(2.2, 44) = 2.1,<br>p = 0.13 | F(4, 76) = 0.771,<br>p = 0.547 |

Note. Significant three-way interaction was found only in Experiment 1. To follow up the significant three-way interaction, we grouped the data by vowel, and analysed the sample two-way interaction between phase and axis. There was no significant simple two-way interaction between phase and axis for any of the vowels (bod: p = 0.055, bad: p = 0.051, bead: p = 0.6). \*p < 0.05 \*\* p < 0.001.

**Figure S2.**
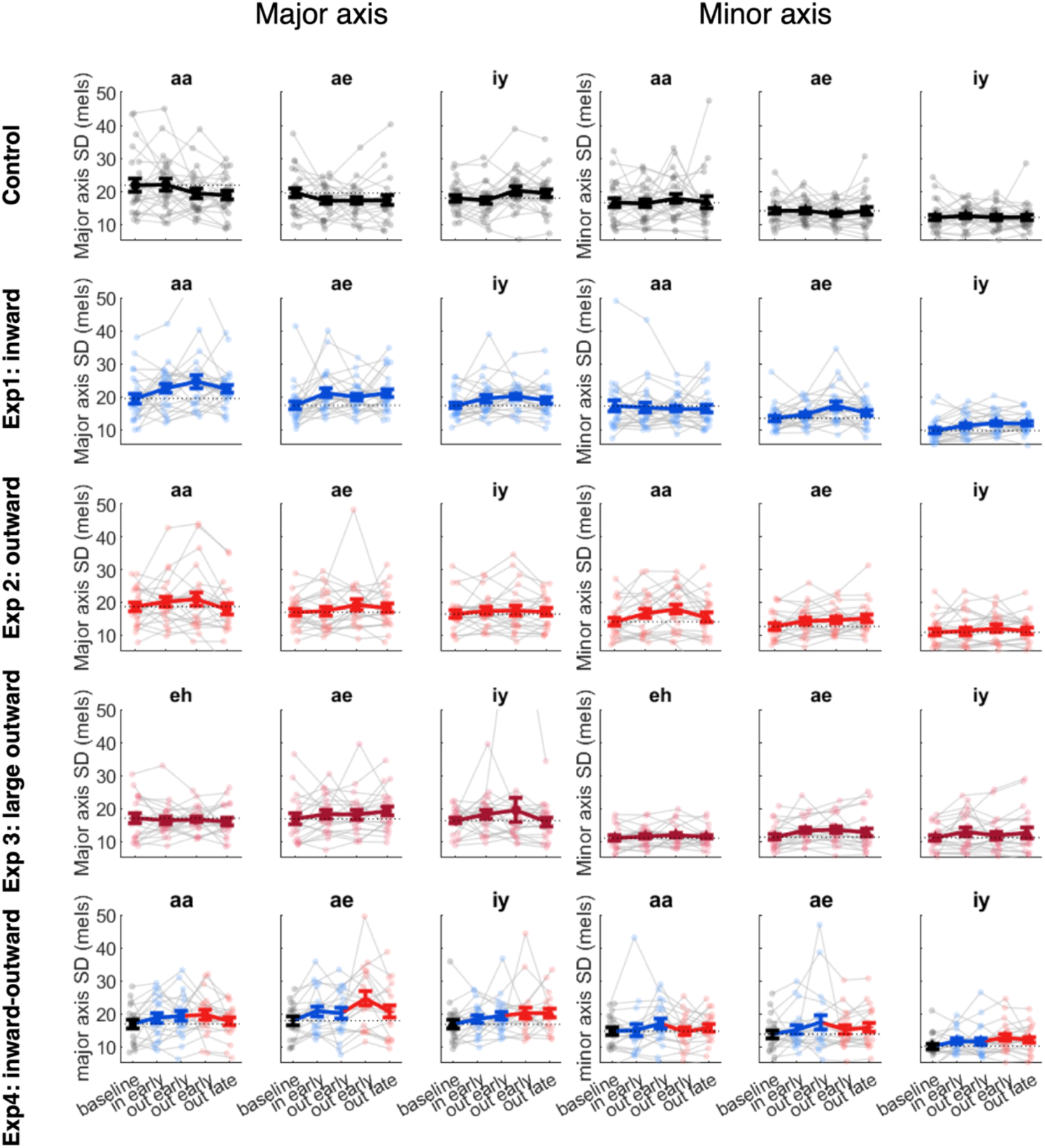
Variability changes along major and minor axes in different words. Individual (small transparent dots, thin lines) and group means (large solid dots, thick lines) of formant variability in the baseline, exposure and washout phases. Error bars show standard error.

**Table S2.**
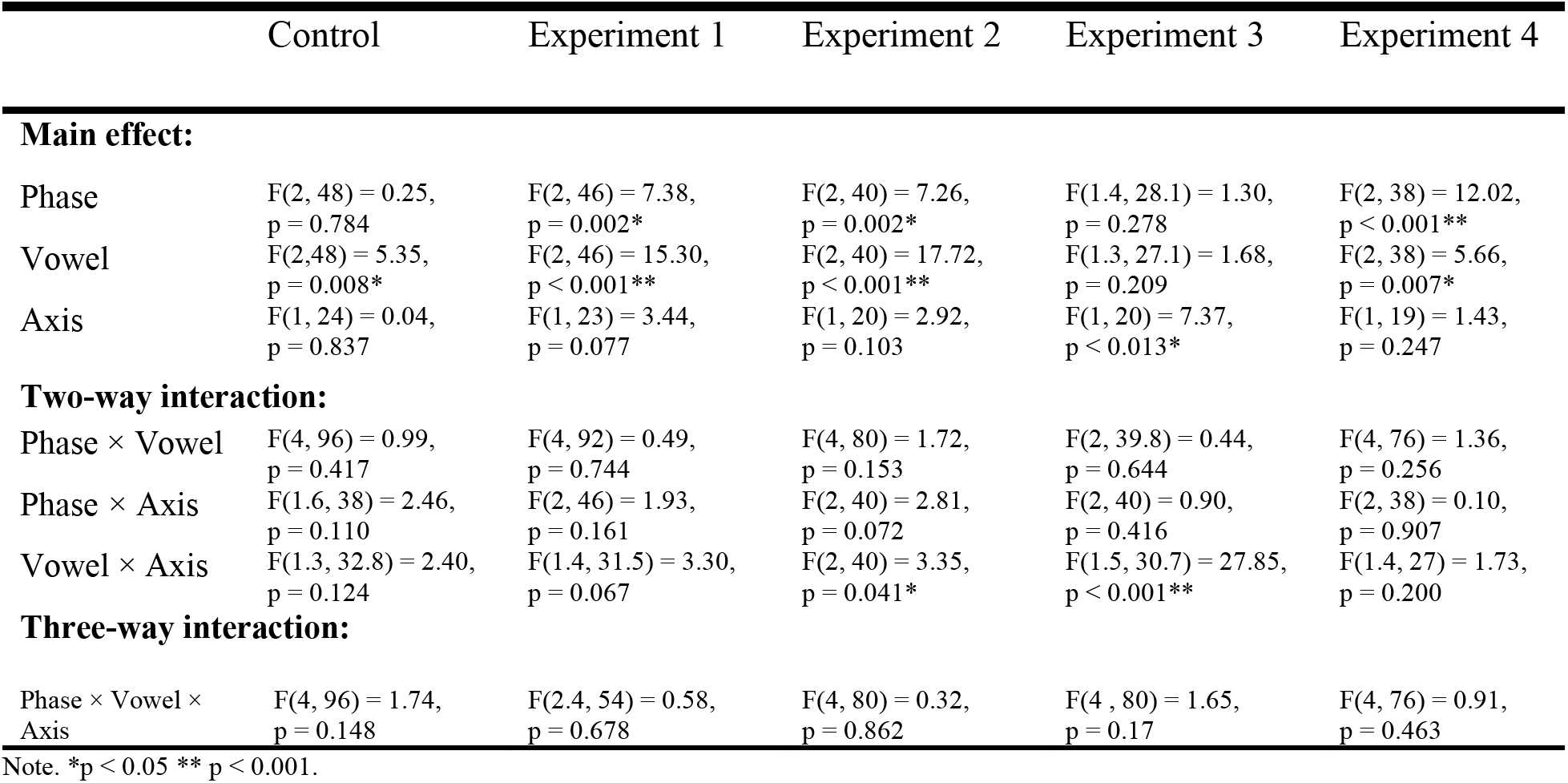
Variability (SD) changes along F1 and F2 axis.

|  | Control | Experiment 1 | Experiment 2 | Experiment 3 | Experiment 4 |
| --- | --- | --- | --- | --- | --- |
| **Main effect:** |  |  |  |  |  |
| Phase | F(2, 48) = 0.25,<br>p = 0.784 | F(2, 46) = 7.38,<br>p = 0.002* | F(2, 40) = 7.26,<br>p = 0.002* | F(1.4, 28.1) = 1.30,<br>p = 0.278 | F(2, 38) = 12.02,<br>p < 0.001** |
| Vowel | F(2, 48) = 5.35,<br>p = 0.008* | F(2, 46) = 15.30,<br>p < 0.001** | F(2, 40) = 17.72,<br>p < 0.001** | F(1.3, 27.1) = 1.68,<br>p = 0.209 | F(2, 38) = 5.66,<br>p = 0.007* |
| Axis | F(1, 24) = 0.04,<br>p = 0.837 | F(1, 23) = 3.44,<br>p = 0.077 | F(1, 20) = 2.92,<br>p = 0.103 | F(1, 20) = 7.37,<br>p < 0.013* | F(1, 19) = 1.43,<br>p = 0.247 |
| **Two-way interaction:** |  |  |  |  |  |
| Phase × Vowel | F(4, 96) = 0.99,<br>p = 0.417 | F(4, 92) = 0.49,<br>p = 0.744 | F(4, 80) = 1.72,<br>p = 0.153 | F(2, 39.8) = 0.44,<br>p = 0.644 | F(4, 76) = 1.36,<br>p = 0.256 |
| Phase × Axis | F(1.6, 38) = 2.46,<br>p = 0.110 | F(2, 46) = 1.93,<br>p = 0.161 | F(2, 40) = 2.81,<br>p = 0.072 | F(2, 40) = 0.90,<br>p = 0.416 | F(2, 38) = 0.10,<br>p = 0.907 |
| Vowel × Axis | F(1.3, 32.8) = 2.40,<br>p = 0.124 | F(1.4, 31.5) = 3.30,<br>p = 0.067 | F(2, 40) = 3.35,<br>p = 0.041* | F(1.5, 30.7) = 27.85,<br>p < 0.001** | F(1.4, 27) = 1.73,<br>p = 0.200 |
| **Three-way interaction:** |  |  |  |  |  |
| Phase × Vowel × Axis | F(4, 96) = 1.74,<br>p = 0.148 | F(2.4, 54) = 0.58,<br>p = 0.678 | F(4, 80) = 0.32,<br>p = 0.862 | F(4, 80) = 1.65,<br>p = 0.17 | F(4, 76) = 0.91,<br>p = 0.463 |

Note. \*p < 0.05 \*\* p < 0.001.

**Figure S3.**
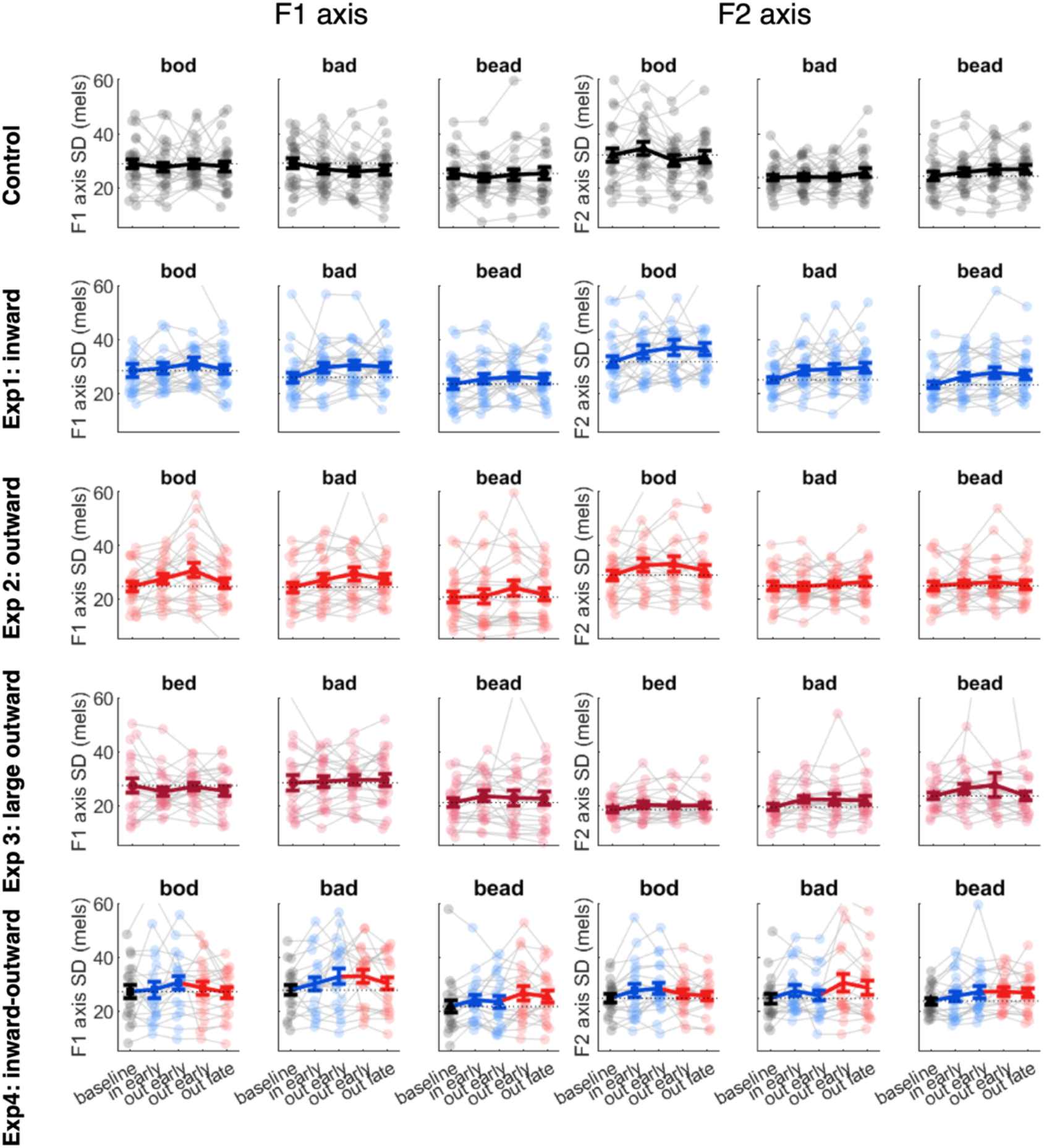
Variability changes along F1 and F2 axes in different words. Individual (small transparent dots, thin lines) and group means (large solid dots, thick lines) of formant variability in the baseline, exposure and washout phases. Error bars show standard error.

## Notes

### Competing Interest Statement

The authors have declared no competing interest.

